# Genome-wide DNA methylation predicts environmentally-driven life history variation in a marine fish

**DOI:** 10.1101/2021.01.28.428603

**Authors:** Hugo Cayuela, Clément Rougeux, Martin Laporte, Claire Mérot, Eric Normandeau, Maëva Leitwein, Yann Dorant, Kim Præbel, Ellen Kenchington, Marie Clément, Pascal Sirois, Louis Bernatchez

## Abstract

The molecular mechanisms underlying intraspecific variation in life history strategies are still poorly understood, despite the importance of this question for understanding of organism’s responses to environmental variability. Theoretical work proposed that epigenetic mechanisms such as DNA methylation might regulate intraspecific variation in life history strategies, however this assumption has rarely been verified empirically in wild populations. We examined associations between genome-wide methylation changes and environmentally-driven life history variation in two lineages of a marine fish that diverged approximatively 2.5 Mya, the capelin (*Mallotus villosus*), from North America and Europe. In both lineages, capelin harbour two contrasted life history strategies: some are strictly semelparous, experience fast actuarial senescence, but benefit from high hatching success by spawning on demersal sites where water temperature is low and relatively stable. In contrast, others are facultative iteroparous, have slower actuarial senescence, and suffer from lower hatching success by breeding in the intertidal zone where temperature is warmer, thermohaline parameters are less stable, and egg desiccation risk is high. Performing whole genome and epigenome sequencing, we showed that these contrasted life history strategies are more likely governed by epigenetic changes than by differences in DNA sequence. While genetic differentiation between the capelin harbouring different life history strategies was negligible, we detected parallel genome-wide methylation changes across lineages. We identified 1,067 differentially methylated regions (DMRs) comprising 15,818 CpGs, with 22% of them located within 5-kb around genes comprising promotor regions. We found that all DMRs were hypermethylated in demersal-spawning individuals. This striking result suggests that lower water temperature at demersal sites leads to an overall hypermethylation of the genome determined during the epigenetic reprogramming occurring over embryonic development. Our study emphasizes that parallel epigenetics changes in lineages with divergent genetic background could have a functional role in the regulation of intraspecific life history variation.

## Introduction

Spatio-temporal variability of the environment is a critical evolutionary driver shaping adaptive strategies in the wild (Blanquart et al. 2013, Savolainen et al. 2013). It determines the schedule of energy allocation of organisms over their lifetime, by affecting the resources that they invest in somatic maintenance (i.e., lifespan), growth, and reproduction (Stearns 1976, 1989). The theory of life history strategies (i.e., covariation between the items of energy expenditure; Promislow & Harvey 1989, Gaillard & Yoccoz 2003) states that the uncertainty of breeding success caused by environmental variability selects for a longer reproductive lifespan and a reduced clutch size at each breeding event (Murphy 1968, Wilbur & Rudolf 2006). Theoretical demographic models showed that extending reproductive lifespan allows organisms to reproduce on multiple occasions and adjust their breeding effort in a flexible way, which permits mitigation of the risk of reproductive failure in variable and unpredictable environments (Bulmer 1985, Tuljapurkar 2013). Field and experimental studies validated those theoretical expectations and showed that the level of environmental variability regulates the reproductive lifespan duration among different populations of the same species (Nevoux et al. 2010; Cayuela et al. 2016, 2019; Lind et al. 2020). However, the molecular mechanisms underlying such intraspecific variation in lifespan are still poorly understood, despite the tremendous importance of this topic for our understanding of organism’s responses to environmental variability.

Our current knowledge of the genetic basis of lifespan and its correlations with other life-history components is currently restricted to a handful of genomic studies that have highlighted the polygenic architecture of these traits, mostly in model organisms (e.g., Browner et al. 2004, Valenzano et al. 2015, Austad & Hoffman 2018, Flatt & Partridge 2018). Although reproductive lifespan is under partial genetic control, both demographic and genetic quantitative studies have generally revealed the low heritability (*h*^2^) and plastic character of life history components under spatiotemporal environmental variation (Price & Schluter 1991, Merilä & Sheldon 1999, Hoffmann et al. 2016). Therefore, one may expect that alternative molecular mechanisms able to modulate gene expression without any change in the DNA sequence (referred to as epigenetics; e.g., Goldberg et al. 2007), might play a central role in the regulation of lifespan and life history strategies (Parrott & Bertucci 2019).

DNA methylation (called methylation hereafter) is the most extensively studied epigenetic mechanism (Smith & Meissner 2013, Klutstein et al. 2016) and has been reported to be a strong predictor of lifespan and aging across mammalians (Pal & Tyler 2016, Lowe et al. 2018) and very recently in fishes (Anastasiadi & Piferrer 2020). For instance, using the methylation status of ~300 cytosines across the genome, one may predict the chronological age of an individual with an error of only 3.6 years (correlation coefficient 0.96) and 3.33 weeks (correlation coefficient 0.84) in human and mouse, respectively (Horvath 2013, Stubbs et al. 2017). Despite the unprecedented accuracy of recently developed epigenetic clocks, the discrepancy between an individual’s methylation age and chronological age varies, and the magnitude and directionality of this discordance are associated with physiological function and life history traits (Parrott & Bertucci 2019). In humans, an acceleration of methylation age correlates with an increase of mortality risk (Marioni et al. 2015), likely because accelerated epigenetic aging is associated with increased risk of age-related diseases (Horvath & Raj 2018) and aberrant gene expression (Reynolds et al. 2014). Although the discrepancies between methylation and chronological ages seem partially determined by the genetic background (Marioni et al. 2015), environmental conditions experienced during early development could exert a strong influence on the degree of epigenetic-to-chronological age discordance experienced later in life (Parrott & Bertucci 2019).

Here, we tested the association between genome-wide methylation (CpG islands) and intraspecific variation of life history strategy associated with contrasted environmental spawning conditions (**Fig.1**). Capelin (*Mallotus villosus*), a small marine pelagic fish with a circumpolar distribution, displays two life history strategies (Christiansen et al. 2008; **Fig.1**) anchored in multiple ancient lineages that diverged approximatively 2.5 Mya (Dodson et al. 2007, Cayuela et al. 2020). Capelin adopting demersal spawning are strictly semelparous (i.e., single reproductive event over lifetime) and experience a strong actuarial senescence (i.e., increase of mortality after breeding) irrespective of sex (Christiansen et al. 2008). They reproduce at spawning sites located at water depths ranging from 10 to 300 m, where environmental conditions are relatively predictable, and water temperature is stable and low (Penton et al. 2012). Fish adopting the beach spawning strategy are facultative iteroparous (i.e., one or two reproductive events over lifetime) in both sexes, suffer a weaker post-breeding senescence, and reproduce in the intertidal zone where water temperature is relatively warm but variable and desiccation risk caused by wind makes offspring survival highly unpredictable (Frank & Leggett 1981, Leggett & Frank 1984, Penton et al. 2012). Previous studies showed negligible genetic variation between pools of breeders of the two life history strategies in the North Atlantic (Præbel et al. 2008, Kenchington et al. 2015, Cayuela et al. 2020).

**Fig.1.**
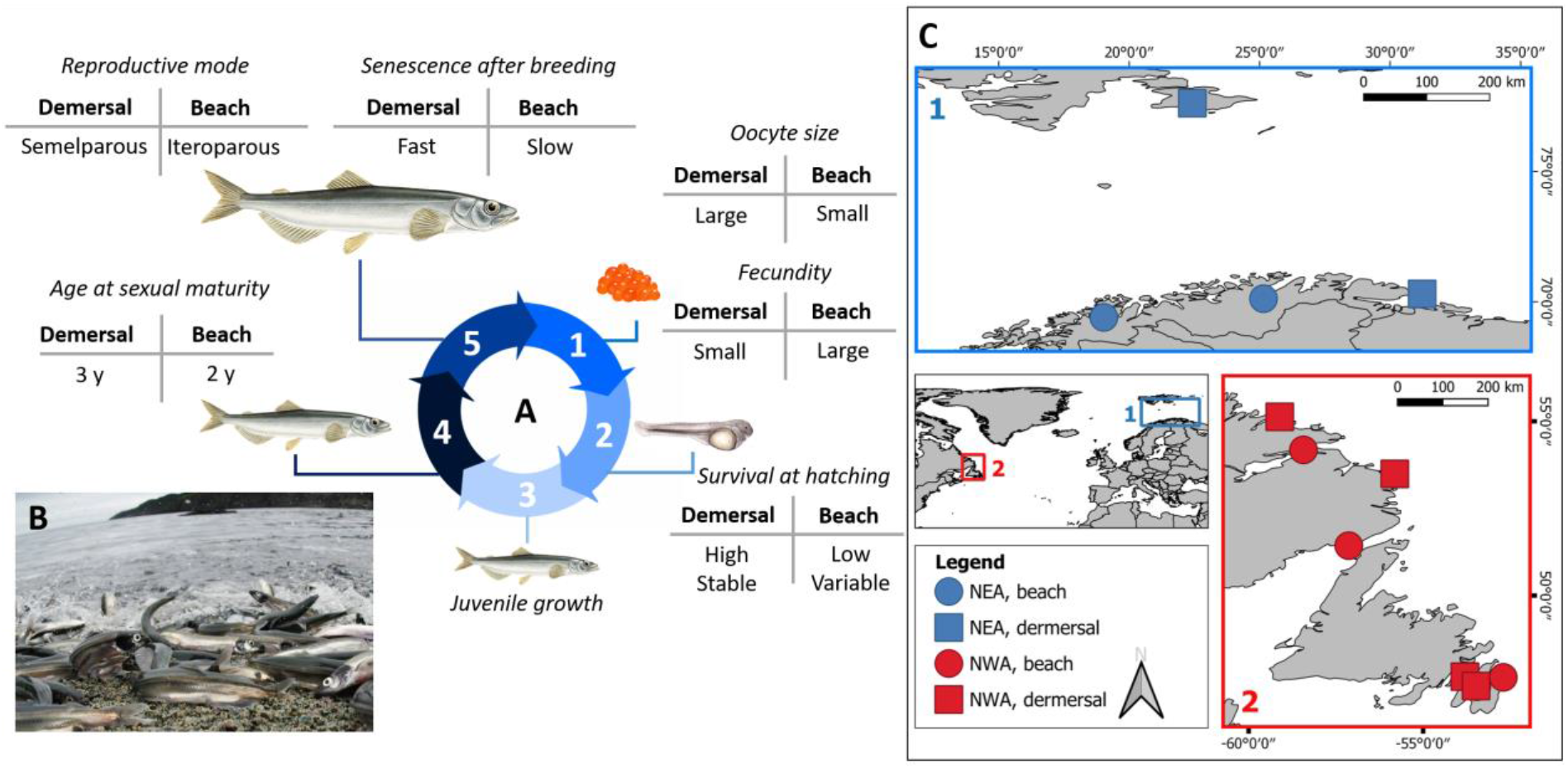
Alternative life history strategies in the capelin (*Mallotus villosus*) based on the work of Christiansen et al. (2008) and study area. (A) Life history differences between demersal-spawning and beach-spawning individuals. (1) Oocyte stage: demersal-spawning individuals reproduce at spawning sites located at water depth ranging from 10 to 250 m where environmental conditions are relatively cold, stable and predictable. By contrast, beach-spawning individuals breed in the intertidal zone (B) where highly variable environmental conditions (i.e., variable but generally warmer temperature and wind) are experienced. Demersal-spawning females produce less eggs than beach-spawning females, but their eggs are larger. Egg development rate is drastically shorter due to warmer temperature in beach-spawning sites than in demersal spawning sites (Penton et al. 2012) (2-3) Larval and juvenile stages: survival at hatching is high and relatively stable over space and time in demersal spawning individuals whereas it has the opposite characteristics in beach spawning individuals (Penton et al. 2012). Larval and juvenile growth are performed at sea and no information about life history is available. (4) Subadult stage: sexual maturity is usually reached at 2 or 3 years in the two life history phenotypes (Sirois, unpublished data). (5) Adult stage: when they are sexually mature, demersal-spawning males and females reproduce once (i.e., strict semelparity) and experience a strong actuarial senescence within the weeks following reproduction. By contrast, beach-spawning individuals, especially females, may reproduce over two successive years (i.e., facultative iteroparity) and suffer a weaker actuarial senescence (i.e. increase of mortality with age). Beach-spawning females does not suffer from fecundity loss during the second reproductive event, suggesting negligible reproductive senescence.

We generated whole-genome-sequencing (WGS) and whole-genome-bisulfite-sequencing (WGBS) data for both demersal and beach-spawning individuals from two ancient lineages occurring in the north-east Atlantic Ocean (NEA lineage) and the north-west Atlantic Ocean (NWA lineage). An association of life-history strategy with patterns of methylation would support a role of epigenetics as a molecular mechanism underlying the intraspecific variation of reproductive lifespan and life history strategy. First, using WGS data, we (1) quantified the extent of genetic divergence between demersal and beach-spawning individuals within each lineage. Then, using WGBS data, we (2) quantified the total amount of methylation independent of the genetic background, and (3) we quantified the total amount of methylation variation associated with lineages and history life strategy. To further support the hypothesis that no genetic basis underlies the association between methylation and history life strategy, we tested for this association after controlling for genetic background. Lastly, controlling for lineage, we identified differentially methylated CpGs (DMLs) and genomic regions (DMRs) associated with life history strategies.

## Methods

### Sample collection and whole genome resequencing data

We analyzed samples of 453 breeding adults (241 and 212 beach spawning and demersal individuals respectively) from 12 spawning sites (6 demersal sites and 6 beach spawning sites; **Fig.1**) located in the northwest (8 sites, NWA lineage) and northeast Atlantic (4 sites, NEA lineage) (for details see (see **Supplementary material S8**, **Table S1**). The DNA samples from two sites (DRL and BB65) were reused from a previous study (Kenchington et al. 2015). The fish were collected and a piece of dorsal fin was preserved in RNAlater or 96% EtOH. DNA was extracted with a salt-based method (Aljanabi & Martinez 1997) and an RNAse A (Qiagen) treatment was applied following the manufacturer’s recommendation. DNA quality was assessed using gel electrophoresis. A total of 453 individuals from the 12 spawning sites were sequenced for the whole-genome (WGS) at low coverage (~1.5X per individual) with the median sample size per site being 38 (range: 20 to 50) individuals. A subset of 55 individuals from 11 spawning sites (excluding the DRL site for which DNA quality was too low for WGS) was sequenced at high coverage (~17X per individual) for whole-genome following a bisulfite conversion steps to tackle methylated cytosines (WGBS) (see **Supplementary material S8**, **Table S1**).

### Molecular analyses: whole-genome sequencing

DNA quality of each extract was evaluated with nanodrop and on a 1% agarose gel electrophoresis. Only samples with acceptable ratios that showed clear high molecular weight bands were retained for library preparation. Following the approach used in Therkildsen & Palumbi (2017), we removed DNA fragments shorter than 1kb by treating each extract with Axygen beads in a 0.4:1 ratio, and eluted the DNA in 10mM Tris-Cl, pH 8.5. We measured DNA concentrations with Biotium Accuclear and normalised all samples at a concentration of 5ng/μL. Then, sample DNA extracts were randomized, distributed in 17 plates (96-well) and re-normalised at 2ng/μL.

Whole-genome high-quality libraries were prepared for each fish sample according to the protocol described in previous studies (Baym et al. 2015, Therkildsen & Palumbi 2017). Briefly, a tagmentation reaction using enzyme from the Nextera kit, which simultaneously fragments the DNA and incorporates partial adapters, was carried out in a 2.5 μl volume with approximately 2 ng of input DNA. Then, we used a two-step PCR procedure with a total of 12 cycles (8+4) to add the remaining Illumina adapter sequence with dual index barcodes and amplify the libraries. The PCR was conducted with the KAPA Library Amplification Kit and custom primers derived from Nextera XT set of barcodes (total 384 combinations – Table SX). Amplification products were purified from primers and size-selected with a two-steps Axygen beads cleaning protocol, first with a ratio 0.5:1, keeping the supernatant (medium and short DNA fragments), second with a ratio 0.75:1, keeping the beads (medium fragments). Final concentration of the librairies were quantified with Biotium Accuclear and fragment size distribution was estimated with an Agilent BioAnalyzer for a subset of 10 to 20 samples per plate. Equimolar amount of 84 librairies were combined into 6 separate ppols for sequencing on 6 lanes of paired-end 150bp reads on Illumina HiSeq2000 system.

Raw reads were trimmed and filtered for quality using the default parameters with FastP (Chen et al. 2018). Reads were aligned to the reference genome with BWA-MEM (Li & Durbin 2009) and filtered with samtools v1.8 (Li et al. 2009) to keep only unpaired, orphaned, and concordantly paired reads with a mapping quality over 10. Duplicate reads were removed with the MarkDuplicates module of Picard Tools v1.119 (http://broadinstitute.github.io/picard/). Then, we realigned reads around indels with the GATK IndelRealigner (McKenna et al. 2010). Finally, to avoid double-counting the sequencing support during SNP calling, we used the clipOverlap program in the bamUtil package v1.0.14 (Breese and Liu, 2013) to soft clip overlapping read ends and we kept only the read with the highest quality score in overlapping regions. This pipeline was inspired by Therkildsen & Palumbi (2017) and is available at https://github.com/enormandeau/wgs_sample_preparation.

### Molecular analyses: genome-wide DNA methylation

Genome-wide DNA methylation maps generated by whole genome shotgun bisulfite sequencing (WGBS). DNA methylation mapping relied on the detection of cytosine / thymidine polymorphisms after bisulfite conversion. Treatment of genomic DNA with sodium bisulfite induces the deamination of unmethylated cytosine bases to uracil, while methylated cytosine bases remain unchanged. Hence, after PCR, unmethylated cytosines are detected as thymidines whereas remaining cytosines indicate cytosine methylation. The tracks display cytosine DNA methylation as percentage of reads with a thymine versus a cytosine (0 - 100%).

Library preparation and sequencing were performed at the Centre d’Expertise et de Services Genome Québec (Montréal,QC, Canada). Regarding library preparation, Genomic DNA was spiked with unmethylated λ DNA and fragmented. Fragments underwent end repair, adenylation of 3’ends, and adaptor ligation. Adaptor ligated DNA was bisulfite-converted followed by amplification by PCR. Library qualities were assessed using the Agilent 2100 BioAnalyzer (Agilent Technologies).

Libraries were sequenced on Illumina HiSeq2000 system. WGBS data processing was carried out as described by Johnson et al. (2012). In short, reads were aligned to the bisulfite converted reference genome using BWA; (i) clonal reads, (ii) reads with low mapping quality score, (iii) reads with mismatches to converted reference, (iv) reads mapping on the forward and reverse strand of the bisulfite converted genome, (v) read pairs not mapped at the expected distance based on library insert size, and (vi) read pairs that mapped in the wrong direction were removed.

Raw whole genome bisulphite sequencing (WGBS) reads were trimmed and cleaned for quality (≥25), error rate (threshold of 0.15) and adaptor sequence using *trim_galore* v0.4.5 (http://www.bioinformatics.babraham.ac.uk/projects/trim_galore/). Trimmed sequences were aligned to the capelin reference genome (Cayuela et al. 2020) using *BSseeker2* v2.1.5 (Guo et al. 2013) with *Bowtie2* v2.1.0 (Langmead & Salzberg 2012) in the end-to-end alignment mode. Duplicates were flagged in BAM files with the *Picard-tools* v1.119 program (http://broadinstitute.github.io/picard/), before determining methylation levels for each site by using the *BSseeker2* methylation call step. The raw methylation file was filtered by removing C-T DNA polymorphism identified from called genotypes, in order to avoid ‘false’ methylation variation at those positions (Le Luyer et al. 2017). We used the *CGmapTools* suite v0.1.1 (Guo et al. 2017) to extract only CpGs sites determined as CG context (avoiding CHH and CHG contexts), and with a coverage bounded between a minimum of 10X and a maximum of 100X in order to avoid noise from repetitive elements and putative paralog genes. Using this set of parameters, we identified a total of 11,073,309 and 11,562,801 methylated loci (CpGs) in the lineage NWA and NEA respectively.

### SNP identification and genome-wide divergence among lineages and life history strategies

Analyses of WGS data were carried out using ANGSD v0.931 (Korneliussen et al., 2014), a software specifically designed to take genotype uncertainty into account instead of basing the analysis on called genotypes. The analytical pipeline is available at https://github.com/clairemerot/angsd_pipeline. We kept input reads with a samtools flag below 255 (not primary, failure and duplicate reads, tag - remove_bads = 1), with a minimum base quality score (minQ) of 20, a minimum mapping quality score (minMapQ) of 30q. We used the GATK genotype likelihood model (GL 2) to generate allelic frequency spectra, infer major and minor alleles (doMajorMinor 1), and estimate allele frequencies (doMaf 2). We filtered to keep only SNPs covered by at least one read in at least 50% of the individuals, with a total coverage below 1812 (4 times the number of individuals) to avoid including repeated regions in the analysis, and with minor allele frequency above 5%. In this way, we identified 6,711,583 SNPs. We then calculated *F_ST_* between lineages and life history strategies, randomly subsampled to a similar size of 59 individuals. We used the realSFS function in ANGSD providing the previously obtained list of polymorphic SNPs and their polarisation as major or minor allele (options – sites and – doMajorMinor 3).

### Decomposition of the genetic and methylation variation associated with lineages and life history strategies

To illustrate patterns of genetic variation, we extracted the genotype likelihood matrix of the 53 individuals (that were also sequenced from WGBS), inferred the individual covariance matrix with *PCAngsd* (Meisner & Albrechtsen 2018) and decomposed it onto orthogonal axes using principal component analysis (PCA) using a scaling 2 transformation, which added an eigenvalue correction, to obtain the individuals PC scores (Legendre & Legendre 1998). For epigenetic variation, PC scores were similarly obtained from the CpGs methylation percentage matrix on the exact same individuals. All graphics representing genetic and epigenetic PC-axes show the average scores for four different groups (2 lineages × 2 habitats) with a 95% confidence interval. To assess if genetic variation can explain epigenetic variation, we produce a backward selection of the genetic PC-axes (*P* < 0.1) on the response epigenetic matrix. All selected genetic PC-axes were then used to produce a redundancy analysis (RDA). The significance of the global model and individual genetic PC-axes were then tested with analyses of variance using 1,000 permutations of the data. The percentage of epigenetic variation explained by genetic variation was then computed based on an adjusted R^2^. Next, we compared how genetic and epigenetic variation can be explained by lineages and life history strategies. Thus, we produced RDAs where lineage and life history strategy variables explained genetic or epigenetic PC-axes. An analysis of variance partitioning was produced on each RDA result to quantify the percentage of variance explained by: i) both lineage and life history strategy (i.e., shared variation), ii) lineage or life history strategy alone, and iii) lineage or life history strategy after controlling for the other variables. All statistics were computed with the software R, using the ‘vegan’ package.

### Genome-wide DMLs and DMRs candidates associated with lineages and life history strategies

We applied a generalized linear model (GLM) implemented in the *DSS* R package (Wu et al. 2013) to identify differentially methylated loci (DMLs) and regions (DMRs) between lineages (NEA vs NWA) using life history strategy as a control covariate. Then, we applied another GLM to identify DMLs and DMRs between life history strategies using lineage as a control covariate. For both GLMs, we controlled for interaction between terms, aiming to compare directly both lineages and reproductive strategies. CpGs showing a probability of *Bonferroni-corrected P* < 0.1 were defined as DMLs. DMRs were retained when at least 10 DMLs occurred in a minimum sequence of 100bp with a probability threshold of *Bonferroni-corrected P* < 0.1. We allowed merging of DMRs less distant than 50bp to be defined as the same DMR. As quantitative metrics to characterize significant DMRs, we used a combination of the statistic describing the area difference under the curve (AreaStat) between methylation levels in DMR between conditions and the mean methylation level per DMR.

## Results

### Assessing genomic variation between lineages and life history strategies

Whole-genome-sequencing analyses showed pronounced genetic variation between NWA and NEA lineages, but negligible genetic differentiation between individuals of the two life history strategies within both lineages. RDA indicated that lineage was associated with 53.9% of the genetic variance (Global significance of both models: *P* < 0.001) whereas no genetic variation associated with life history strategy was observed (**Fig.2**). Pairwise *F_ST_* estimated between lineages and life history strategies confirmed this pattern (Table 1): *F_ST_* was 0.23 between lineages and about two orders of magnitude less between life history within lineages (NWA: 0.003; NEA: 0.004).

**Fig.2.**
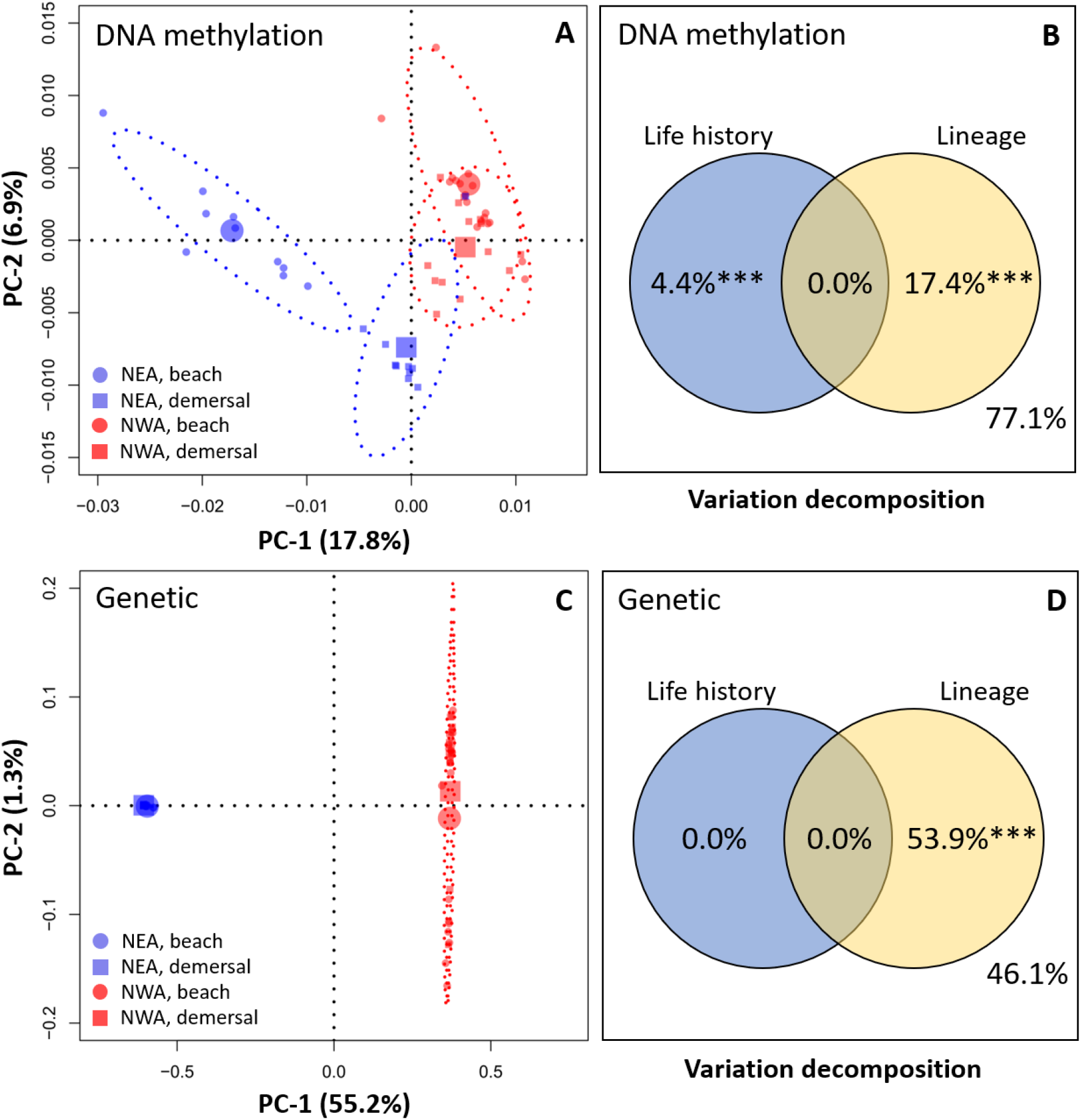
Principal component analyses (A and C) and partial redundancy analyses (B and D) showing the proposition of methylation (A and B) and genetic (C and D) variation associated with life history (beach spawning *vs* demersal individuals) and lineages (NEA and NWA). In A and C: centroids represent the mean of the four groups (NEA beach, NEA demersal, NWA beach, and NWA demersal) and ellipses show 95% confidence intervals. In B and C: the proportion of variation non-explained by the model (i.e., residual variance) is 77.1% (B) and 46.1% (C).

**Table 1.** pairwise *F_ST_* between linages (NWA and NEA) and life history strategies (beach-spawning individuals BS, and demersal individuals D) within both lineages. Note that 95% CI cannot be calculated in ANGSD and are therefore not provided.

|  | NWA_BS | NWA_D | NEA_BS | NEA_D |
| --- | --- | --- | --- | --- |
| NWA_BS | - | - | - | - |
| NWA_D | 0.0032 | - | - | - |
| NEA_BS | 0.2250 | 0.2254 | - | - |
| NEA_D | 0.2369 | 0.2373 | 0.0042 | - |

### Quantifying methylation variation between lineages and life history strategies

Genome-wide methylation differed importantly between lineages (17.4% of variation explained by the lineage, P < 0.001). In contrast with genomic variation, the RDA showed that life history strategy also explained a significant proportion (4.4%, P < 0.001) of the total methylome variation (**Fig.2**). Our analyses also showed that a substantial part of the methylation variation associated with life history strategy was independent from the genetic background. Thus, we first quantified the amount of methylation that was explained by the genetic background as a whole. After backward selection, seven genetic PC-axes (PC-1, 35, 41, 42, 45, 48 and 52) representing 57.9% of genetic variation were kept (*P* < 0.1) for the redundancy analysis to explain methylation variation. A total of 28.2% of the genome-wide methylation variation was significantly explained by genetic variation (Global significance of the model: *P* < 0.001; only PC-1 and 35 where highly significant *P* < 0.001, the others PC-axes were significant under a threshold of *P* < 0.1). Then, we built a RDA model where methylation variation was associated with life history strategy after controlling for genetic background. A total of 1.8% of all methylation variation (not explained by genetic background) was still associated with life history strategy (Global significance of the model: *P* < 0.001).

### Identifying differentially methylated regions associated with lineages and life history strategies

Using GLMs, we determined the genomic regions involved in the 17.4% of the methylome varying between lineages. We identified 557,789 and 7,765 significant DMLs and DMRs (**Supplementary material S1),** respectively. The DMRs contained 121,424 CpGs. Seventy-nine percent of the DMRs (6,156 DMRs) were hypermethylated in NEA, indicating an overall hypermethylation trend in this lineage. Nine percent of the 7,765 DMRs (i.e., 739; **Supplementary material S2**) were located within 5-kb region around genes whereas less than 1% of those DMRs (50 DMRs; **Supplementary material S2**) were located within protein-coding genes involved in various molecular and cellular processes (transcript names and GOs provided in **Supplementary material S3**).

We identified 251,442 significant DMLs (**Supplementary material S4**) and 1,067 significant DMRs (**Supplementary material S5**) predicting individual’s life history strategy (**Fig. 3B** and **3D**), and shared in parallel by both lineages despite their pronounced genetic and methylation divergence. The parallel DMRs contained 15,818 CpGs. The unsupervised clustering of DMR methylation profiles first separated both lineages (**Fig 3C**) and then clustered “demersal” life history strategy capelin from each lineage (**Fig 3D**). All (100%) of the 1,067 parallel were hypermethylated in capelin of the demersal life history strategy (**Fig.3A** and **3B**). The DMR were homogeneously spread along the genome (**Fig.3A**): 22% of the DMRs (237 DMRs; **Supplementary material S6**) were located within 5-kb around genes whereas 1% of the DMRs (11 DMRs; **Supplementary material S6**) were located within the sequence of protein-coding genes (transcript names and GOs provided in **Supplementary material S7**). Several of these genes were involved in the regulation of brain and central nervous system development (transcript names: XM_012822573.1; XM_012835203.1; XM_012824747.1; XM_012829347.1; XM_012823759.1) and the immune system (transcript names: XM_012831717.1; XM_012825256.1). The profile of the DMRs between life history strategies showed variable length (**Supplementary material S8**, **Fig. S1A**) and density of CpGs (**Fig. S1B**), with a strong correlation between the number of CpGs per DMR and the score of differentiation (AreaStat) between life history strategies (Pearson’s r=0.55, *P*<0.001, **Fig. S1C**).

**Fig.3.**
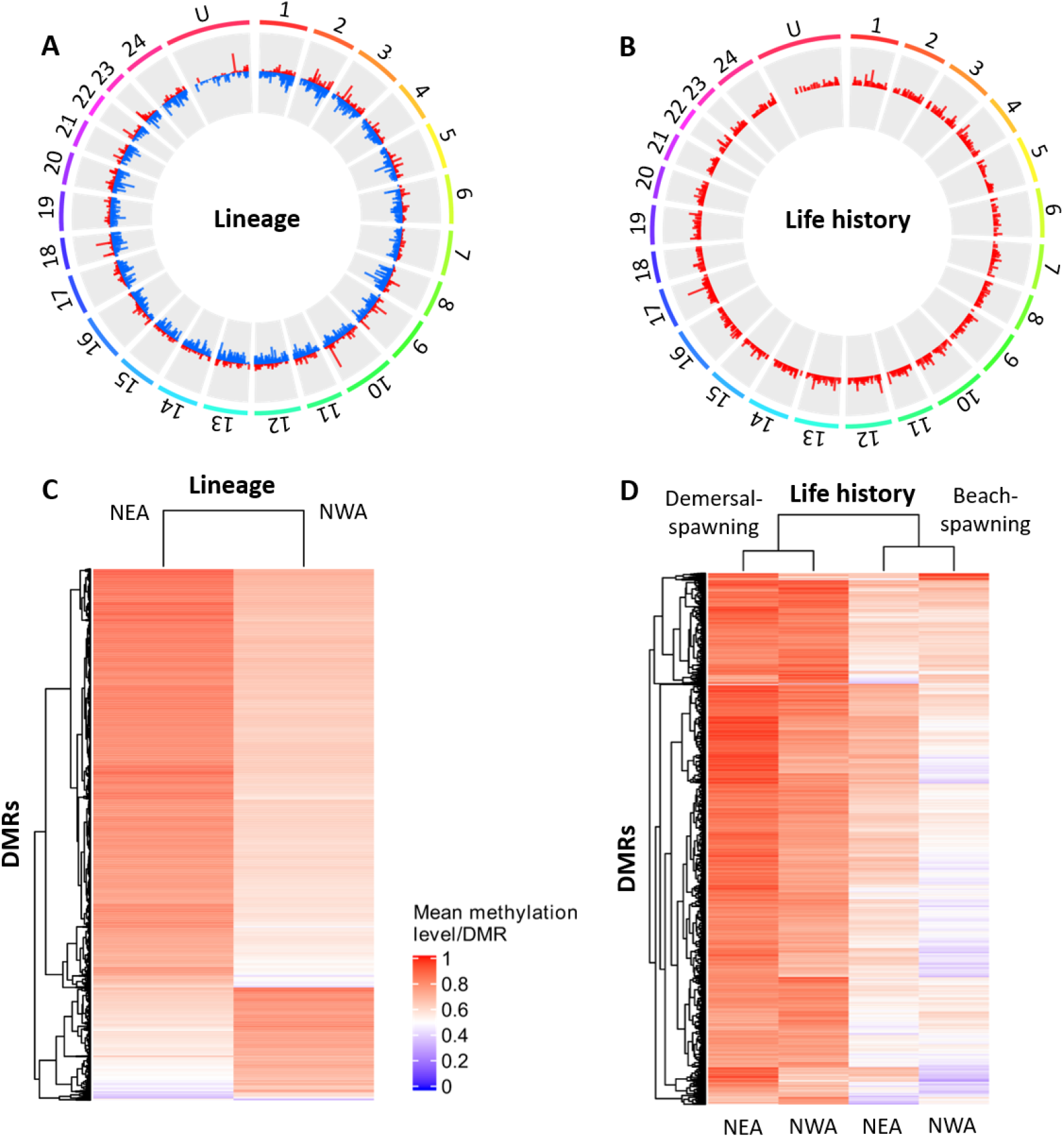
Difference in methylation profiles between lineages (NAE and NWA) and life history strategies (semelparous demersal-spawning individuals vs iteroparous beach-spawning individuals) within lineages. (A-B) Circosplots show the distribution of DMRs associated with lineage and life history across the 24 chromosomes of the capelin. (C-D) Heatmaps showing the mean DNA methylation level of CpGs within identified significant DMRs in both lineages and life history comparisons. Identified DMRs allowed segregating NEA and NWA lineages (C) and clustered similar demersal-spawning and beach-spawning strategies from different lineages (D).

## Discussion

The molecular basis of reproductive lifespan and its correlated life history components are still poorly understood, despite the tremendous importance of this issue to better predict organism’s responses to environmental variability. Our results support the hypothesis that environmentally-related variation in reproductive strategy and related lifespan are more likely governed by epigenetic changes than by differences in DNA sequence in the Capelin. We found negligible genetic variation between the two life history strategies within each capelin lineage from two continents. By contrast, our results showed a strong differentiation in the methylome of demersal- and beach-spawning capelin, supporting a functional role of epigenetics in the regulation of life history strategies in wild populations. Moreover, the methylation differentiation was observed across the two capelin lineages that diverged approximately 2.5 Mya (Dodson et al. 2007, Cayuela et al. 2020), demonstrating that similar methylome modifications are associated with the differential expression of life history strategies that evolved on both sides of the North Atlantic.

### Methylation variation between ancient capelin lineages

Our results show that, after controlling for life history strategy, 17% of genome-wide methylation variation is associated with ancient lineages that began to diverge approximatively 2.5 MyA. Factors responsible for this level of divergence are speculative at this time but inter-lineage differences could originate in the divergent environmental characteristics of capelin marine habitat in the NEA and NWA, respectively. Indeed, it has been shown that biotic (e.g., trophic resources; Morán et al. 2013) and abiotic factors (e.g., temperature; Campos et al. 2013, Metzger & Schulte 2017) may impact methylation patterns in both marine and freshwater fishes. Furthermore, difference in methylation between lineages might also result from the strong inter-lineage genetic divergence, since 28% of the overall methylation variation is associated with the genetic background in the capelin. A similar effect of the genetic background on methylation has previously been reported in plants (Seymour et al. 2014) and humans (Bell et al. 2011).

### Negligible genetic differentiation between life history strategies

Our study revealed weak genetic difference between capelin of the two life history strategies and this was repeated within the two lineages. The *F_ST_* values (close to 0) indicated pronounced gene flow, if not panmixia, among individuals with the two life history strategies. Moreover, the amount of genetic variation associated with life history strategy was negligible. This result is congruent with previous studies that have revealed marginal genetic variation between beach and demersal spawning sites within the NWA lineage using mitochondrial DNA, microsatellites and SNPs (Dodson et al. 1991, Præbel et al. 2008, Kenchington et al. 2015, Cayuela et al. 2020). Nevertheless, the absence of marked genomic differentiation between demersal-spawners and beach-spawners does not rule out the possibility of a partial limited control of these life history strategies through a limited number of genes. Indeed, Cayuela et al. (2020) identified 105 SNP outliers associated with capelin life history strategies in the lineage NWA, suggesting that genetic variation between demersal-spawners and beach-spawners is likely maintained in some genomic regions via spatially varying selection and/or habitat matching choice despite high gene flow.

### Differences in methylation between life histories strategies across different lineages

Our results revealed that 4% of the methylation variation is associated with life histories after controlling for lineages and that 251,442 DML and 1,067 DMRs were implicated in this differentiation. Moreover, we showed that after controlling for the whole genetic background, 1.8% of the methylation variation associated to life history strategy difference was shared between both lineages. This indicates parallel environmentally induced variation of methylation across both capelin lineages and supports the role of epigenetics as one of the molecular mechanisms involved in the intraspecific shifts of life history strategy. Strikingly, the 1,067 DMRs recurrently presented higher level of methylation in demersal-spawning individuals than in beach-spawning individuals in both capelin lineages.

The differences in methylation associated with life history strategies are likely induced by water temperature experienced during embryonic development, the stage at which temperature usually influences most DNA methylation patterns in fishes (Anastasiadi et al. 2017, Burgerhout et al. 2017, Lallias et al. 2020, Sävilammi et al. 2020). This interpretation is supported by an experimental study on *Dicentrarchus labrax* (Anastasiadi et al. 2017), which found that genome-wide reprogramming events of methylation marks induced by temperature usually take place shortly after fertilization and are completed during embryogenesis rather than later in life (i.e., larvae and juvenile stages). It is also noteworthy that epigenomic modifications mediated by temperature are very unlikely to occur in adults when they return to reproduce, since they usually spend less than 24 hours on spawning sites (Davoren 2013). Moreover, embryonic development at low temperature is expected to result in the genome-wide hypermethylation observed in our study. A comparative study across fish species showed that cold water is usually associated with a general increase of methylation level along the genome (Varriale & Bernardi 2006; but also see Metzger & Schulte 2017). Penton et al. (2012) reported higher hourly and mean daily incubation temperatures at a beach spawning site, relative to demersal spawning site, on the northeast Newfoundland coast, which was supported by a literature review of temperatures at spawning locations in the wider North Atlantic. Therefore, it is plausible that differences in thermal conditions prevailing during embryonic growth in beach- and demersal-spawning sites may be responsible for the observed methylation differences in adult capelin. This association supposes that adults return preferentially reproduce in their habitat of birth (i.e., habitat matching choice), but not necessarily at the same locations unlike the homing behavior observed in salmonid fishes for instance (Keefer & Caudill 2014). Habitat matching choice (*sensu* Edelaar et al. 2008) has previously been proposed as a mechanism to explain local adaptation to environmental conditions prevailing in capelin beaching-spawning sites despite high gene flow (Cayuela et al. 2020).

Our results showed that hypermethylation is associated with reduced reproductive lifespan and high actuarial senescence in demersal-spawning individuals. This association suggests that methylation modification induced at the embryonic stage could cause differences in mortality patterns later in life. Indeed, the hypermethylation of regulatory regions or sequence of genes involved in the regulation of central nervous system and immunity responses could lead to stronger aging and shorter reproductive lifespan in demersal-spawning individuals than in beach-spawning ones. Although the existence of such a mechanism in fishes is still unknown, it has been repeatedly observed in humans where a number of hypermethylated genes are associated with age-related diseases, increased senescence, and reduced lifespan (Post et al. 1999, So et al. 2006, Zampieri et al. 2015). The overall hypomethylation of those genes in beach-spawning individuals could contribute to longer lifespan and slower actuarial senescence, thus allowing facultative iteroparity. If iteroparity effectively allows increasing individuals fitness in variable environments as expected by theoretical models (Bulmer 1985, Tuljapurkar 2013), methylation modifications experienced at early stage would be adaptive and could provide a selective advantage to beach-born individuals reproducing in their natal highly variable habitat. Our study would thus provide empirical evidence for parallel adaptive plasticity allowing fish from lineages that have divergence for 2.5 Mya to cope with similar sources of environmental variability. Admittedly, further experimental studies will be needed to rigorously test this hypothesis and examine how methylation and gene expression are associated with lifespan and aging rate of demersal- and beach-spawning individuals in common garden experiments.

### Research avenues

Our study supports the hypothesis that thermal conditions experienced during early development may affect genome-wide methylation patterns later in life and indicated that those epigenetic changes are associated with the expression of very distinct life history strategies. As such, this study offers novel research avenues, notably pertaining to the epigenetic clock which generality and properties have poorly been investigated so far in ectotherms (but see Anastasiadi & Piferrer 2020 for fish). It also raises important questions about the influence of thermal variation induced by climate change on the epigenome of ectotherms, and their consequences on individual life trajectories and ultimately the demography of wild populations.

## Acknowledgements

We thank biologists and technicians of the Department of Fisheries and Oceans Canada for their implication as well as all everyone who contributed to sampling throughout the study area. This research was funded by a Strategic Project Grant from the Natural Sciences and Engineering Research Council of Canada (NSERC) to L. Bernatchez, M. Clément and P. Sirois, a financial contribution of Resources Aquatiques Québec and was also supported by in-kind contribution from many other organisations: Department of Fisheries and Oceans Canada, Nunatsiavut Government, NunatuKavut Community Council, Labrador Fishermen’s Union Shrimp Company, Department of Fisheries and Aquaculture – Government of Newfoundland and Labrador, World Wildlife Fund Canada, St. Lawrence Global Observatory, Parc Marin du Saguenay–Saint-Laurent, Manitoba University, and the Greenland Institute of Natural Resources. Hugo Cayuela was supported by a Vanier-Banting postdoctoral fellowship and the Swiss National Science Foundation (grant number: 31003A_182265).

## Contribution statement

H.C. wrote the paper. H.C., C.R., C.M., M.L., and E.N. made the bioinformatics and the statistical analyses. L.B., M.C. and P.S. initiated the project, and L.B. conceptualized and coordinated the work. All authors read and edited the final manuscript version.

## Notes

### Competing Interest Statement

The authors have declared no competing interest.

