## Supplementary_material for "Genome-wide DNA methylation predicts environmentally-driven life history variation in a marine fish": Supplementary_material_S8.pdf

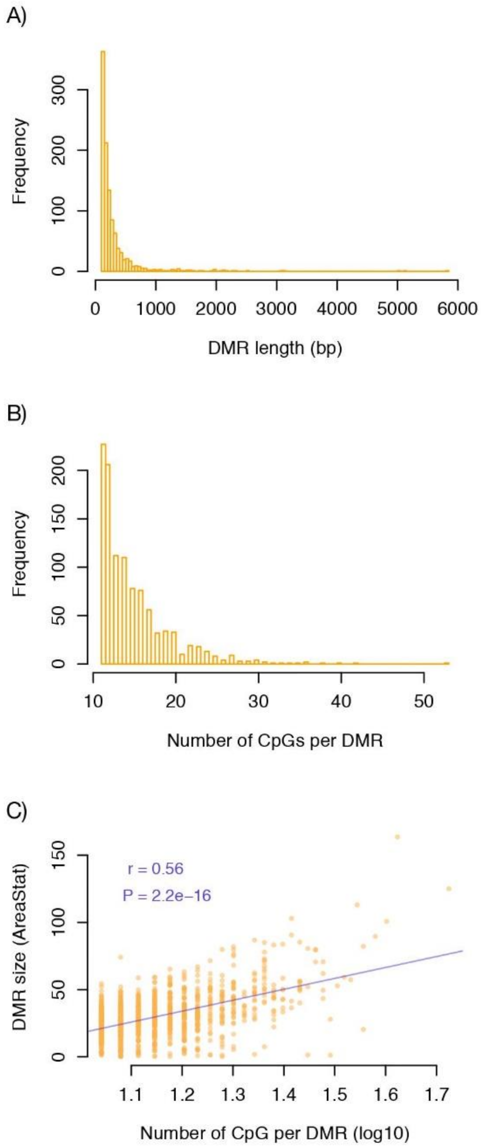


**Fig.S1.** DMR size and length and CpG content. (A) Frequency of DMRs according to their length (in bp). (B) Frequency of DMRs according to their content of CpG (number of CpGs). (C) DMR size (AreaStat) according to their content of CpG (log10(number of CpGs)).

**Table S1**. Spawning sites in the two glacial lineages of capelin (NEA and NWA).

| ID | Latitude | Longitude | Sample_size_WGS | Sample_size_WGBS | Lineage | Life history |
|---|---|---|---|---|---|---|
| BAL | 69.4 | 19.033333 | 39 | 5 | NEA | Beach spawner |
| BSO | 70.3 | 31.233333 | 39 | 5 | NEA | Dermersal spawner |
| BSW | 77.61666 | 22.416666 | 40 | 5 | NEA | Dermersal spawner |
| POR | 70.11666 | 25.15 | 20 | 5 | NEA | Beach spawner |
| BB65 | 47.67299 | -53.802196 | 49 | 5 | NWA | Dermersal spawner |
| BEL-B | 47.38165 | -53.46462 | 40 | 5 | NWA | Beach spawner |
| BEL-D | 47.40355 | -53.48096 | 36 | 5 | NWA | Dermersal spawner |
| BLA | 51.4264 | -57.1313 | 46 | 5 | NWA | Beach spawner |
| DRL | 53.493333 | -55.813333 | 28 | 0 | NWA | Dermersal spawner |
| MAK2 | 55.12708 | -59.1085 | 20 | 5 | NWA | Dermersal spawner |
| MID | 47.650694 | -52.696313 | 50 | 5 | NWA | Beach spawner |
| RIG | 54.1799 | -58.4288 | 46 | 5 | NWA | Beach spawner |
